# Neurostructural and cognitive signatures of novel polygenic risk scores for molecular brain aging

**DOI:** 10.1101/2024.11.17.624029

**Authors:** Kevan P. Clifford, Fernanda Dos Santos, Mohamed Abdelhack, Daniel Felsky, Etienne Sibille, Yuliya S. Nikolova

## Abstract

The world population is shifting sharply toward an older-age demographic. To navigate the escalating burden of physical and cognitive decline common to aging, and heightened risk of neurodegenerative and neuropsychiatric disease, we require advances in treatment and prevention interventions. These advances are predicated on attaining a deeper understanding of the molecular processes underlying brain aging. Here, we employed novel GWAS and cis-eQTL-based polygenic risk scores (^GWAS^AGE-PRS and ^cis-eQTL^AGE-PRS) indexing genetic risk for accelerated molecular brain aging, and examined their associations with cortical thickness and performance in age-sensitive cognitive domains in 31 384 participants (16 392 women, age 64.1±7.65) from the UK Biobank. While ^GWAS^AGE-PRS was nominally associated with lower cortical thickness in frontotemporal regions, ^cis-eQTL^AGE-PRS displayed robust associations with greater cortical thickness in age-sensitive frontal, temporal, and parietal regions, including the left and right precentral (pFDR<0.0001, pFDR=0.05), left insula (pFDR=0.05), as well as the right supramarginal (pFDR=0.05) and precuneus (pFDR=0.05) regions. Similar pFDR trending associations occurred bilaterally in the caudal middle frontal (pFDR=0.052, pFDR=0.078) and right insula (pFDR=0.071). These structural findings co-occurred alongside increased executive function performance on the Trail Making Test B (pFDR=0.035), suggesting a potential neurostructural and cognitive reserve phenotype. This resilience profile may reflect previously uncharacterized pathways of brain reserve in age-related pathology, informing future translational research identifying novel treatment and prevention targets.

## 1. Introduction

The world population is shifting toward a disproportionately large older-age demographic, with the number of individuals aged 65 and over projected to rise to 22.4% by 2100 (Gu et al., 2021). Given the anticipated burden of age-related cognitive deficits and risk of neurodegenerative and neuropsychiatric diseases, elucidating the molecular processes underlying brain aging is imperative for identifying novel treatment and preventative interventions towards healthy brain aging.

While aging-related changes in the human brain were once thought to be generalized and non-specific, more recent studies have demonstrated conserved age-related processes in distinct molecular and cellular pathways in the brain (Bishop et al., 2010; Erraji-Benchekroun et al., 2005). Notably, normal brain aging has been associated with morphological changes in neuronal structure, particularly reduced complexity of dendritic arborization (Dickstein et al., 2007), diminished synaptic plasticity (Bergado and Almaguer, 2002), altered glial cell structure and function (Torres and Cardenas, 2020), immune system shifts toward neuroinflammation (Di Benedetto et al., 2017), as well as accumulated DNA damage (Maynard et al., 2015). Normal aging also brings gradual cognitive decline, most pronounced in the domains of episodic and working memory, attention, processing speed, fluid intelligence, as well as executive functioning, which is particularly age-sensitive (Buckner, 2004; Li et al., 2001).

Interestingly, inter-individual variability in brain aging trajectories exist, with disparity occurring between individuals’ chronological age and estimated biological brain age. Conceptually, accelerated brain aging refers to neurobiological aging processes occurring more rapidly compared to chronological age (Christman et al., 2020). Estimates of brain age have been derived from cortical neuroimaging (Franke and Gaser, 2019) and epigenetic clocks (Marioni et al., 2019), with the difference between estimated brain age and chronological age reflected by a ‘delta’ value. Extensive prior studies suggest that structural and epigenetic aging may be accelerated in mental illness (e.g., depression and schizophrenia (Han et al., 2021; Koutsouleris et al., 2014), and neurodegenerative conditions (Beheshti et al., 2019). However, the precise molecular mechanisms underlying accelerated brain aging, and their potential application to treatment and prevention of age-related pathology, remain incompletely characterized (Isaev et al., 2018). Further, the cross-sectional nature of much of the published work in this area (Higgins-Chen et al., 2021; Mishra et al., 2023) makes it challenging to determine whether accelerated brain aging is a cause or consequence of the observed conditions.

Currently, assessing molecular brain aging is only possible through postmortem brain tissue sampling, coupled with multi-omics approaches. Brain tissue transcriptomics comprehensively quantify mRNA expression levels, and this intermediary between gene and protein offers insight into both environmental and genetic contributions to complex disease-related traits (Bahn et al., 2001). Cortical gene expression follows developmental trajectories shaping neocortex organization, and is thought to potentially influence structural breakdown of cortical architecture in middle age and late age (Fjell et al., 2015; Huffman, 2012). In previous work (Lin et al., 2020), we utilized postmortem tissue from two age-sensitive prefrontal cortex regions to identify highly age-related genes with robust monotonically increasing or decreasing relationships between gene expression and age (20-90 years). By leveraging the gene expression trajectories of the age-related genes, we developed a construct for individual-level prediction of molecular brain age. We found that in postmortem brain tissue from clinical samples, high delta molecular brain age (molecular brain age - chronological age) was strongly associated with diagnoses of schizophrenia (SCZ) and bipolar disorder (BD) (Lin et al., 2020).

To further bridge these insights to the living human brain, we developed novel indices of genetic risk for accelerated molecular brain aging through two distinct polygenic risk score (PRS) approaches. One approach identified risk variants based on their cis-eQTL status for individual age-dependent genes (^cis-eQTL^AGE-PRS), while the other identified non-overlapping risk variants through a genome-wide association study (GWAS) of whole-transcriptome molecular brain age (^GWAS^AGE-PRS). Across methods (cis-eQTL and GWAS) and thresholds, the PRSs were variably associated with schizophrenia (SCZ) and major depressive disorder (MDD) diagnosis, as well as lower cognitive performance. ^cis-eQTL^AGE-PRSs were additionally associated with Alzheimer’s disease (AD) diagnosis, and postmortem measures of neuritic plaques and neurofibrillary tangles (Lin et al., 2020). These proof-of-principle studies suggest that accelerated molecular brain aging may be at least partially genetically driven. This, in turn, raises the possibility that AGE-PRSs may have more subtle neurostructural and cognitive impacts, detectable in non-clinical populations in the absence of, or ahead of, clinically relevant cognitive impairment. Studies using intermediate phenotypes more proximal to disease biology, and more sensitive to normal aging across mid- to late-life (e.g., cortical thickness (van Erp et al., 2018; Frangou et al., 2022; Márquez and Yassa, 2019; Truong et al., 2013), are particularly likely to yield valuable insight into early stage etiology of brain-aging pathologies (Dohm-Hansen et al., 2024) and identify neural and cognitive targets for early intervention.

To that end, here we seek to characterize the phenotypic effects of ^cis-eQTL^AGE-PRS and ^GWAS^AGE-PRS on brain structure and cognition in a population-representative sample of middle-aged and older adult participants from the UK Biobank. We hypothesized that AGE-PRS will be associated with lower cortical thickness above and beyond the effects of chronological age, and that this effect will be most pronounced in frontal, temporal, and parietal regions that are most susceptible to normal age-related thinning (Fjell et al., 2009; Lemaitre et al., 2012; Thambisetty et al., 2010). In addition, we hypothesized associations between AGE-PRS and poorer performance on cognitive tasks probing age-sensitive domains such as executive function.

## 2. Materials and Methods

### 2.1 UK Biobank data

#### 2.1.1 UK-Biobank study population

The UK Biobank is a large ongoing population-representative cohort study consisting of just over 500 000 participants from the United Kingdom, recruited between 2006-2010. The study has collected extensive genotypic and phenotypic data, including genome-wide genotyping, multimodal imaging, questionnaires, physical measures, and accelerometry (Sudlow et al., 2015). For the purpose of this research, we leveraged genome-wide, structural neuroimaging, and cognitive performance data. Polygenic risk scores were computed in 425 631 of 502 413 participants. Related individuals were accounted for using a kinship coefficient of 0.20, reducing participants to 407 050, followed by retaining only those with both structural neuroimaging data and computed PRSs. This resulted in 31 384 individuals (ages 46-82, mean age of 64.1±7.65, F=16 392).

#### 2.1.2 Genome wide data

Briefly, DNA was extracted from blood samples. Whole-genome genotyping was performed using two similar assays. A smaller subset of participants was genotyped at 807 411 markers using the Applied Biosystems UK BiLEVE Axiom Array by Affymetrix, while other participants were genotyped using the closely related Applied Biosystems UK Biobank Axiom Array (825 927 markers) that shares 95% of marker content with the UK BiLEVE Axiom Array. Both arrays capture genome-wide genetic variations (single nucleotide polymorphism (SNPs) and short insertions and deletions (indels)). To increase the number of testable variants, genotypes were imputed from the Haplotype Reference Consortium (HRC) and UK10K haplotype resource. A detailed account of genetic resources and processes of the UK Biobank can be found elsewhere (Bycroft et al., 2018).

### 2.2 Neuroimaging

#### 2.2.1 Neuroimaging protocol

The UK Biobank brain MRI protocol was performed using a 3 Tesla Siemens Skyra scanner (Siemens Healthineers, Erlangen, Germany) with VD13 software and a 32-channel head coil. The present study leveraged T1-weighted images, acquired using a 3D magnetisation-prepared rapid-acquisition gradient-echo (MPRAGE) sequence at 1 mm isotropic resolution. To facilitate reproducibility, identical equipment and software were used across three assessment centres (Newcastle, Cheadle, Reading) without major updates throughout the study. Complete scanner protocol details can be found at http://biobank.ctsu.ox.ac.uk/crystal/refer.cgi?id=2367

#### 2.2.2 Processing of T1-weighted scans

An automated processing pipeline for brain image analysis and quality control was established for the UK Biobank at the University of Oxford’s Wellcome Centre for Integrative Neuroimaging (WIN/FMRIB). The pipeline is primarily based around FSL (FMRIB’s Software Library) (Jenkinson et al., 2012) and other packages, including FreeSurfer (Fischl, 2012). After brain images were obtained at imaging centres, automated quality control identified artefact and equipment issues. Images were then reconstructed from *k*-space on scanner computers and saved in DICOM file format before conversion to NIFTI file format. Pre-processing included removal of faces for participant anonymity, brain extraction, as well as linear alignment and nonlinear warping to a standard MNI152 brain template. For our purposes, we utilized only T1-weighted images, where estimated cortical gray matter was used to automatically generate “imaging-derived phenotypes” (IDPs) summaries of major tissue types. Processing pipelines are described in further detail elsewhere (Alfaro-Almagro et al., 2018; Miller et al., 2016). In this current study, we leveraged Freesurfer-derived IDP tabular output (data field 20227) from the UK Biobank to derive cortical thickness (CT) values for the 62 regions of the Deskin Killiany-Tourville atlas parcellation (Alexander et al., 2019).

### 2.3 Cognitive performance data

Cognitive performance data probing age-sensitive domains were collected at the same visit as the MRI scan (instance 2). We accessed data for six cognitive tests probing age-sensitive domains, including Trail Making Test A (reversed scored) (n=21 263), Trail Making Test B (reverse scored) (n=21 263), Symbol Digit Substitution (n=21 062), Tower Rearranging Task (n=20 872), Numeric Memory (n=21 542), and Fluid Intelligence (n=28,989), measuring processing speed, executive function, processing speed, executive function, working memory, and verbal and numerical reasoning, respectively. Additionally, a seventh measure (Trail Making Test B-A, n=21 263) was created by subtracting Trail Making Test A times from Trail Making Test B times, which is thought to dissociate effects of processing speed from executive function performance (Varjacic et al., 2018). Further details on cognitive tests in the UK Biobank can be found elsewhere (Fawns-Ritchie and Deary, 2020) and at https://biobank.ndph.ox.ac.uk/showcase/label.cgi?id=100026.

### 2.4 Developing polygenic risk scores for molecular brain aging

Further details on the development of the PRSs can be found in our previously published work (Lin et al., 2020), and a high level overview of their development and application in the current study are depicted in Figure 1.

**Figure 1.**
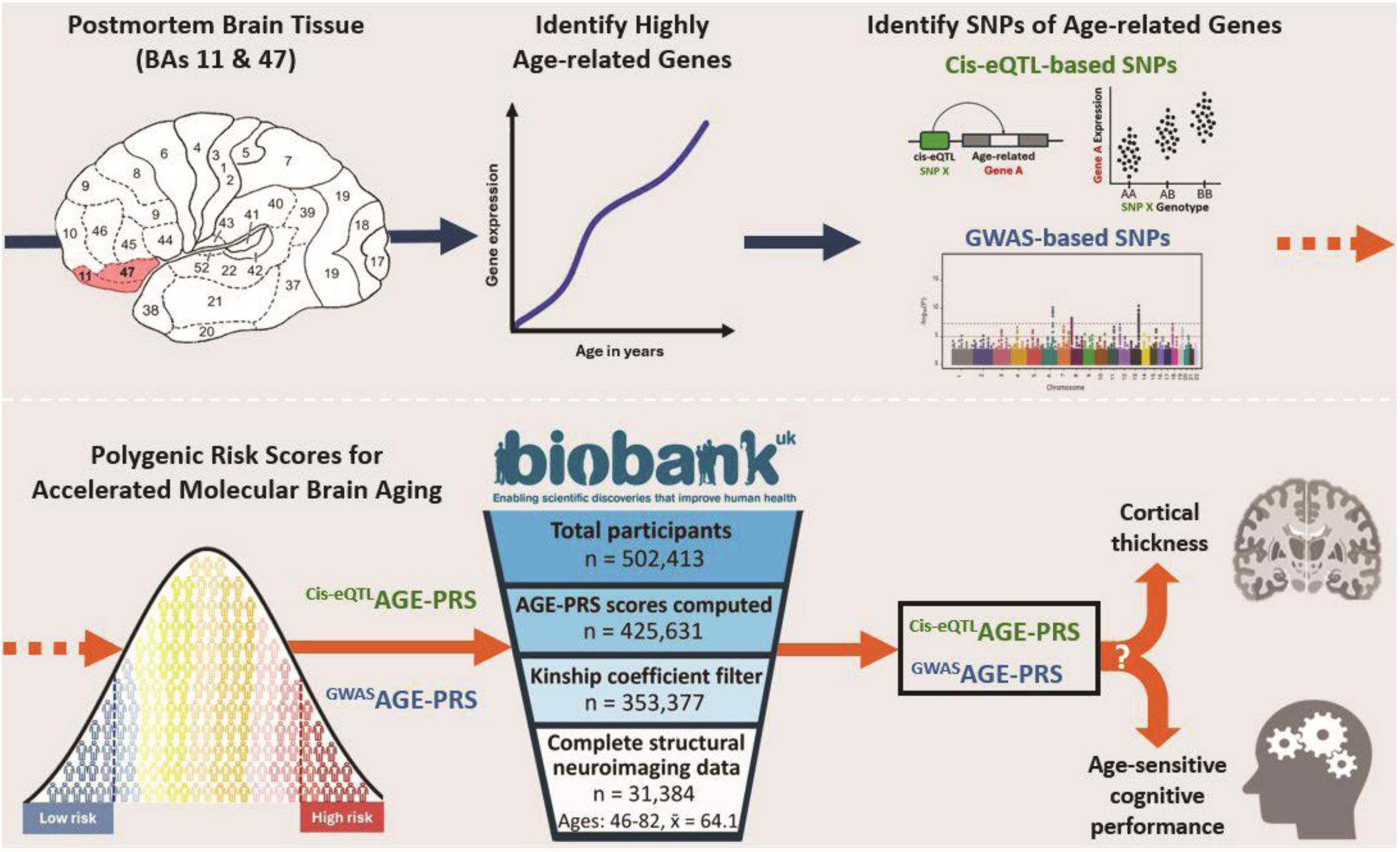
High-level overview of methods. In our previous work (blue arrows) we identified highly age-related genes and leveraged them to identify GWAS and cis-eQTL-based SNPs which were compiled into polygenic risk scores (AGE-PRSs) indexing accelerated molecular brain aging risk. In our current work (orange arrows), we calculated both GWAS based (^GWAS^AGE-PRS) and cis-eQTL-based (^cis-eQTL^AGE-PRS) PRSs in the large, middle age to early old-age UK Biobank cohort. The associations of AGE-PRS with cortical thickness and cognitive performance were evaluated in the UK Biobank.

#### 2.4.1 Identifying SNPs associated with molecular brain age

We obtained gene lists and SNPs identified in our previous work (Lin et al., 2020). Briefly, using postmortem tissue from Brodmann’s areas 11 and 47, highly age-related genes with significant linear relationships between gene expression and age (20-90yrs) were identified. Next, SNPs were identified using two complementary strategies: First, a cis-eQTL analysis was performed, identifying SNPs within 50kb of age-related genes significantly associated with altered gene expression levels consistent with old-age trajectories. Second, a model to predict individual molecular brain age was constructed, leveraging expression trajectories of the age-related genes. A GWAS was then performed, identifying SNPs associated with high delta molecular brain age in each brain region. Cis-eQTL based SNPs were first thresholded by number of identified age-related genes (FDR=10^-4^, 10^-3^, 10^-2^, 5^-2^, 10^-1^) resulting in five groups (Group A=1065 genes, Group B=1548 genes, Group C=2550 genes, Group D=3682 genes, Group E=4425 genes), and further thresholded by cis-eQTL association (q<0.01, q<0.05, q<0.1, q<0.2, q<0.3), resulting in 25 inter-related sets of SNP groups (SNP groups A1:5-E1:5), comprising 94-1836 SNPs. GWAS-based SNPs of interest were identified under different statistical thresholds (p<10^−8^, p<10^−7^, p<10^−6^, p<10^−5^, p<10^-4^), termed SNP groups G1, G2, G3, G4, G5, comprising 3-248 SNPs respectively. SNPs were also tested for associations in independent cohorts, extracted, and aggregated with equal weights at the same thresholds, forming SNP groups G6, G7, G8, G9, G10, comprising 14-394 SNPs. Full details of cis-eQTL and GWAS-derived SNP groups are depicted in Figure 2.

**Figure 2.**
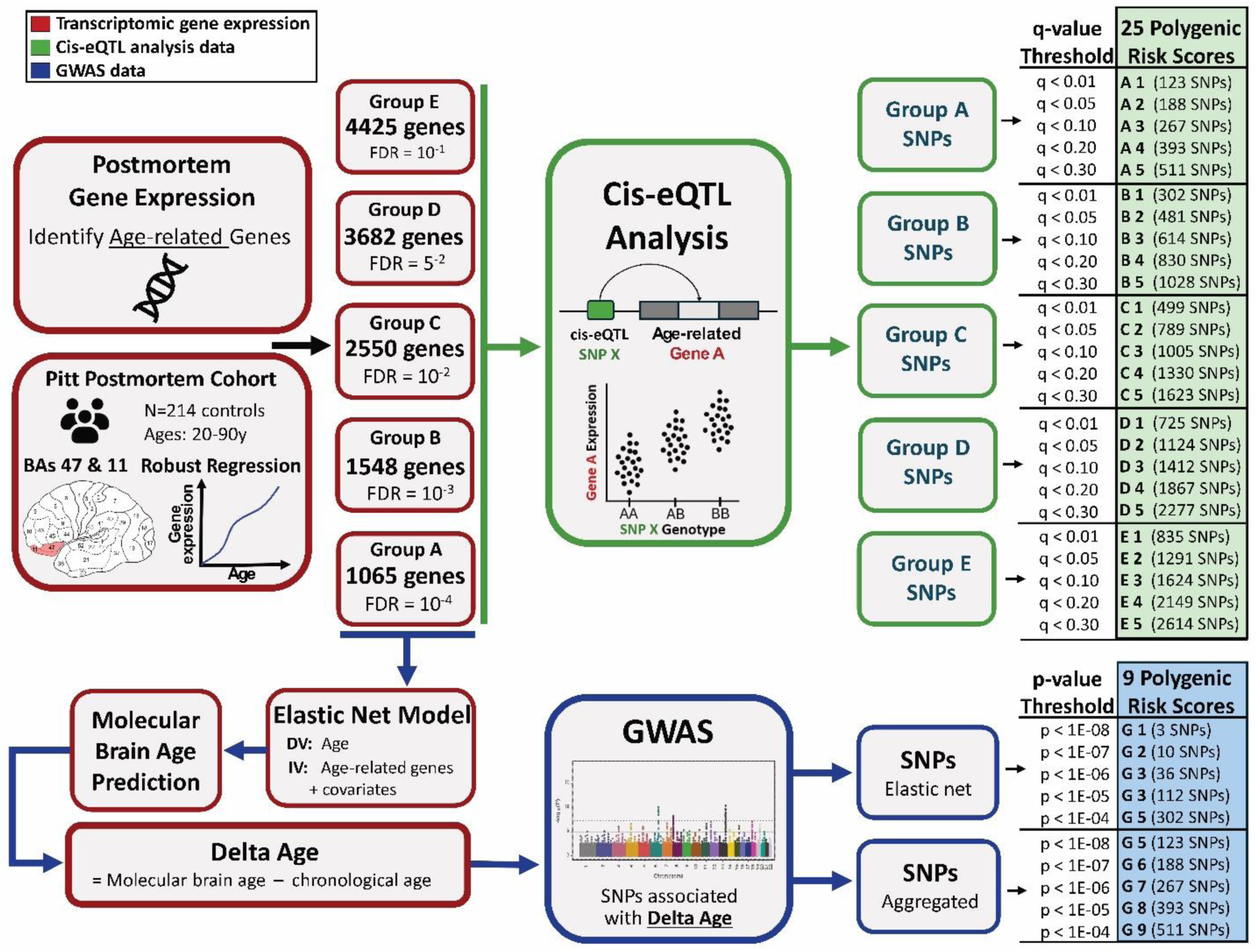
Construction of ^cis-eQTL^AGE-PRSs and ^GWAS^AGE-PRSs. Postmortem tissue from Brodmann areas 47 and 11 (related prefrontal regions) yielded transcriptomic gene expression data. Robust regression was performed between genes’ expression levels and chronological age (ages 20-90) identifying 5 groups of highly age-related genes, based on FDR multiple correction thresholds (A). Cis-eQTL analysis was performed separately for each group of genes, identifying SNPs nearby age-related genes that push gene expression further toward old-age-like trajectories. SNPs were further thresholded by q-values based on association between cis-eQTLs and age, resulting in 25 interrelated polygenic risk scores (B). Separately, expression patterns of the smallest group of age-related genes (Group A, 1065 genes) were used in an elastic net model to predict an individual’s molecular brain age, and resulting delta brain age (molecular brain age – chronological age). A GWAS was performed identifying SNPs associated with delta brain age, thresholded by p-value associations, resulting in 10 interrelated PRSs (C).

A wider range of thresholds were employed than in our prior work (Lin et al., 2020), in order to identify greater numbers of phenotypically relevant SNPs and capture broader patterns of association across thresholds.

#### 2.4.2 Summary statistics

Lastly, summary statistics of SNPs were compiled in our previous work (Lin et al., 2020) with SNPs coded as 0, 1, or 2 in relation to the number of minor alleles. For cis-eQTL-based SNPs, the weight of a SNP was determined by the combination (i.e., multiplication) of the cis-eQTL effect sign and the sign of the age-related gene effect. Accelerated aging was defined as cis-eQTL effects increasing expression of a gene whose expression increases with aging (1 and 1), or cis-eQTL effects decreasing expression of a gene whose expression decreases with aging (-1 and -1). Conversely, delayed aging was defined as cis-eQTL effects decreasing expression of a gene whose expression increases with aging (-1 and 1), or cis-eQTL effects increasing expression of a gene whose expression decreases with aging (1 and -1) . For GWAS-based SNPs, a weight of 1 or -1 was assigned to each SNP based on the sign of the GWAS effect (1 representing accelerated brain aging, -1 representing delayed brain aging).

#### 2.4.3 Construction of polygenic risk scores

In our present work, we leveraged genome-wide data from 502 413 individuals in the UK Biobank. Quality control (QC) steps already performed by the UK Biobank included a missing genotype rate of 0.1, Hardy-Weinberg equilibrium of 1e-15, mach-r2-filter imputation quality of 0.8, and a minor allele frequency of 0.01, all within a European-only population. In addition to these steps, variants with one or more character allele codes were excluded and individual chromosomes were merged using PLINK v1.94 (Purcell et al., 2007). Genetic relatedness was accounted for by retaining only individuals from the ‘used in calculating genetic principal components’ data field (22020)—a proxy for applying a kinship coefficient of 0.20 while retaining one of each pair of related individuals.

After quality control, the summary statistics for cis-eQTL-based SNPs and GWAS-based SNPs were used to calculate individual scores using a method of clumping and thresholding implemented in PLINK1.9 (score function). 25 ^cis-eQTL^AGE-PRS were computed from SNP groups A1:A5, B1:B5, C1:C5, D1:D5, E1:E5, and 10 ^GWAS^AGE-PRS were computed from SNP groups G1:G10. PRSs were subsequently z-score normalized using the scale() function in R.

### 2.5. Statistical Analyses

#### 2.5.1.1 AGE-PRS associations with cortical thickness

All analyses were conducted using R version 4.0.3 (Ihaka and Gentleman, 1996). For each AGE-PRS, linear regressions were conducted with AGE-PRS as the independent variable and regional CT as the dependent variable. Covariates included sex, age, age^2^, (age x sex), (age^2^ x sex), the first 10 genetic principal components, and assessment site. An FDR of 5% was applied for the 62 regional comparisons.

#### 2.5.2 AGE-PRS associations with cognitive performance

For each AGE-PRS, linear regressions were conducted with AGE-PRS as the independent variable and cognitive task scores as the dependent variable. Covariates included sex, age, age^2^, (age x sex), (age^2^ x sex), the first 10 genetic principal components (data field 22009), and assessment site. An FDR of 5% was applied for the 62 regional comparisons.

### 2.6 AGE-PRSs by sex interactions

As an exploratory analysis, linear regressions were conducted for each AGE-PRS, with AGE-PRS as the independent variable, sex as an interaction variable, and regional CT as the dependent variable. Covariates included sex, age, age^2^, (age x sex), (age^2^ x sex), the first 10 genetic principal components, and assessment site. An FDR of 5% was applied for the 62 regional comparisons.

## 3. Results

### 3.1 AGE-PRS associations with cortical thickness

#### 3.1.1 ^cis-eQTL^AGE-PRS associations with cortical thickness

The strongest phenotypic effects between ^cis-eQTL^AGE-PRSs and CT occurred in scores derived from the group D age-related genes (FDR<5^2^, 2550 genes). In the D2 score (q<0.05, 819 SNPs), significantly greater CT was present in the left precentral (pFDR<0.0001) and insula (pFDR=0.05), as well as the right precentral (pFDR=0.05), precuneus (pFDR=0.05), and supramarginal (pFDR=0.05) gyri. Trending levels of greater CT associations occurred bilaterally in both caudal middle frontal gyri (pFDR=0.052, pFDR=0.078) and the right insula (pFDR=0.071). Furthermore, the D1 score (q<0.01, 542 cis-eQTL SNPs) was associated with significantly greater CT in the left precentral (pFDR<0.0001) and insula (pFDR=0.031), as well as the right precentral (pFDR=0.041). The robustness of these regional effects was demonstrated by continuous series of nominal, trending, and significant effects across multiple thresholds (i.e., horizontally in Fig.3). Additional associations occurred in scores from group E of age-related genes (FDR<10^-1^, 4425 genes), particularly the E2 score (q<0.01, 951 cis-eQTL SNPs) where ^cis-eQTL^AGE-PRS was associated with greater CT in the left precentral (pFDR<0.0001) and at trending levels in both left and right caudal middle frontal gyri (pFDR=0.093, pFDR=0.062).

**Figure 3.**
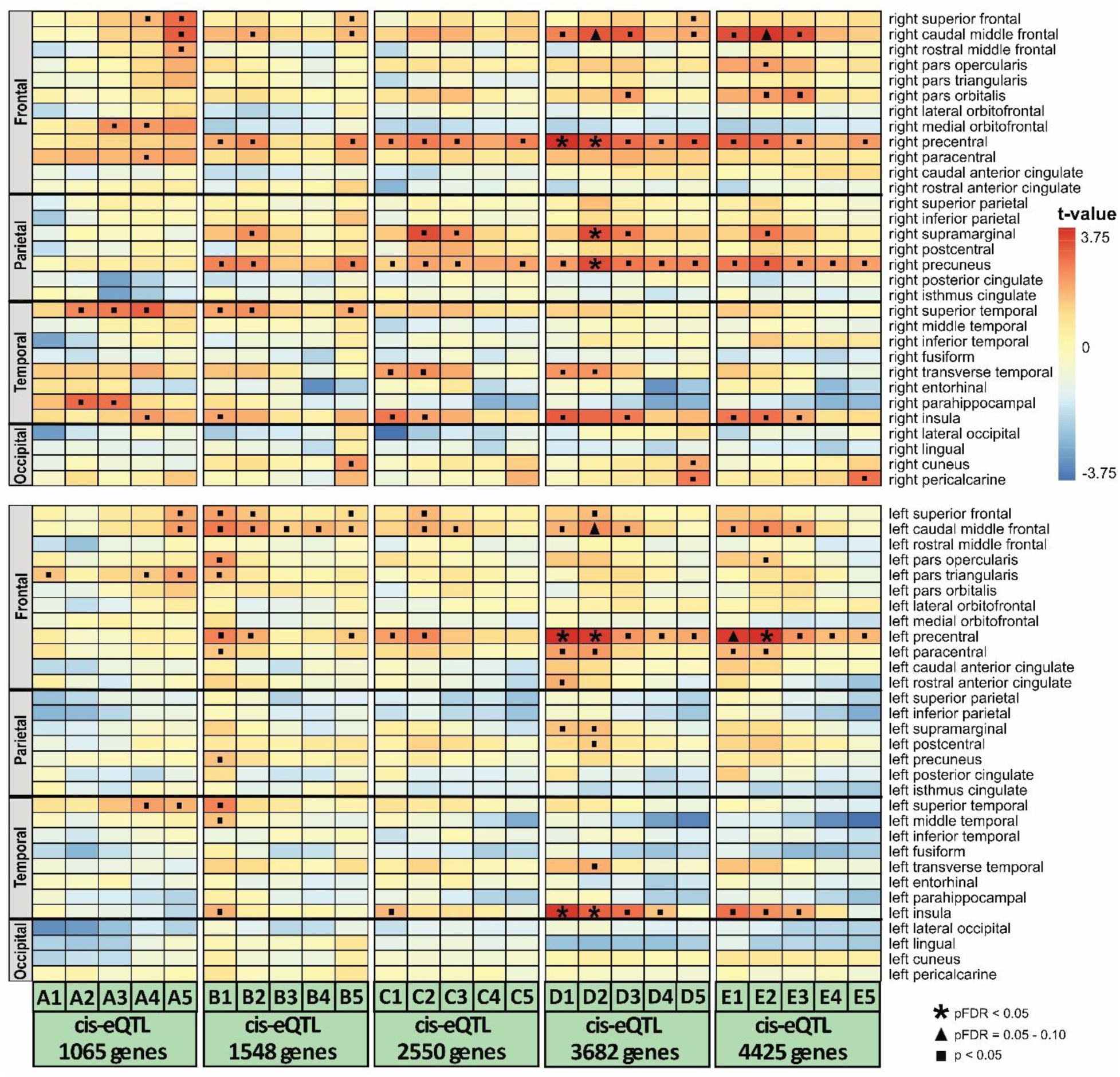
Overview of ^cis-eQTL^AGE-PRSs associations with cortical thickness. Each column corresponds to one of the 25 ^cis-eQTL^AGE-PRSs, while rows depict their respective associations with hemispheric brain regions (top = right hemisphere, bottom = left hemisphere). Within each hemisphere, regions are further organized by frontal, parietal, temporal, and occipital lobes. Warm colors (red/orange) reflect positive associations (i.e., higher PRS → greater cortical thickness), while cool colors (blue) reflect negative associations (i.e., higher PRS → lower cortical thickness) in the respective brain region. Asterisk symbols represent FDR-significant regional associations, while triangle symbols represent FDR-trending associations, and square symbols represent significant associations at p<0.05 uncorrected.

Lastly, a similar association was present at trending level in the E1 score (q<0.01, 630 SNPs) in the left precentral gyrus (pFDR=0.062). Figure 3 summarizes associations across all 25 ^cis-eQTL^AGE-PRSs and CT.

A distributed pattern of nominal associations with greater CT that did not survive FDR correction (p<0.05) occurred across the 25 scores, most frequently in the superior frontal (n=11 ^cis-eQTL^AGE-PRSs), superior temporal (n=9 ^cis-eQTL^AGE-PRSs), paracentral (n=7 ^cis-eQTL^AGE-PRSs), and transverse temporal (n=5 ^cis-eQTL^AGE-PRSs) gyri. Figure 4 depicts regional ^cis-eQTL^AGE-PRS and CT associations. Full statistical results can be found in Supplementary File 1.

**Figure 4.**
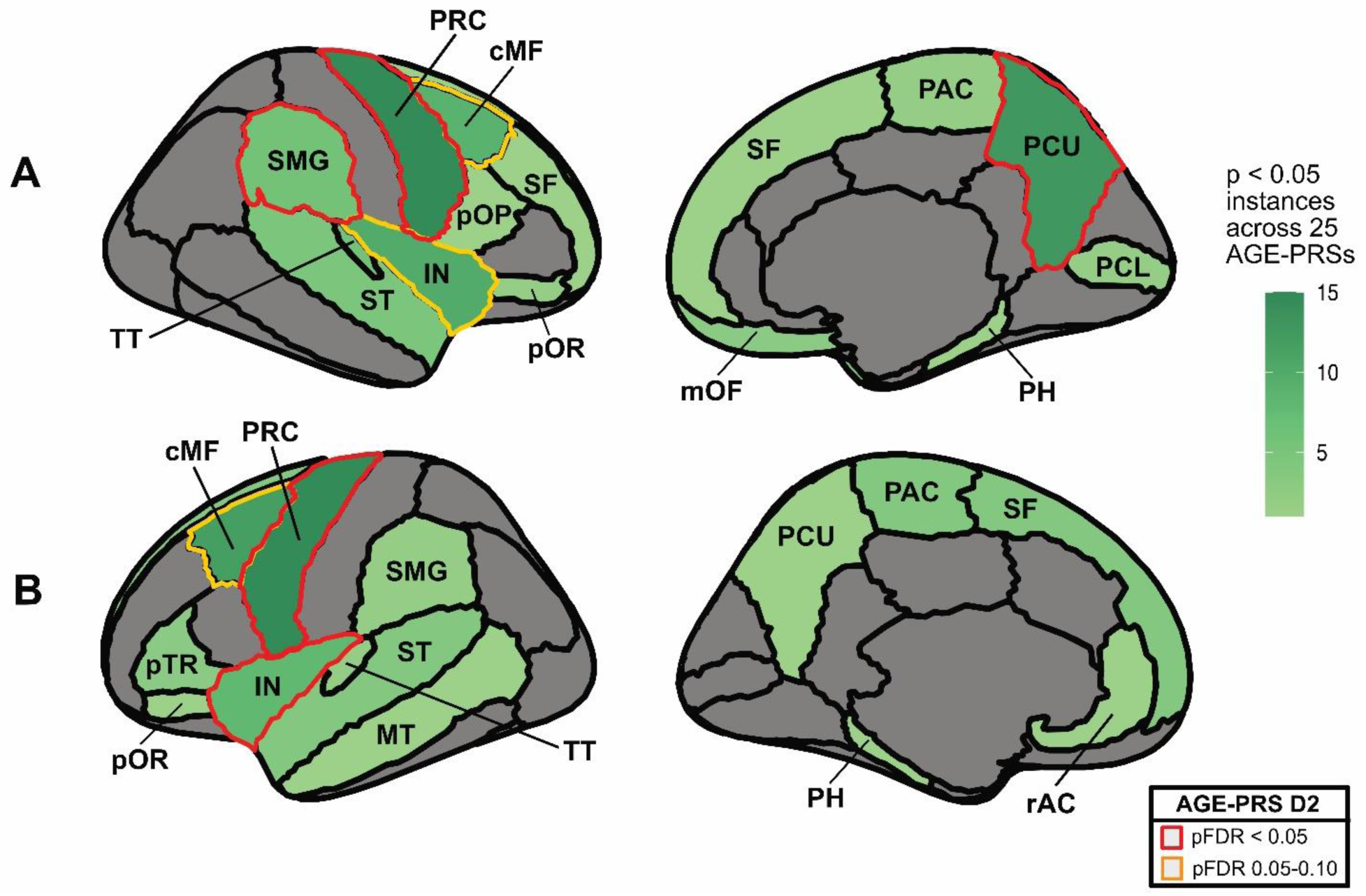
Regional brain map of ^cis-eQTL^AGE-PRSs associations with cortical thickness across thresholds. Brain atlases depicting greater cortical thickness (green) are displayed for the right hemisphere (A) and left hemisphere (B). Darker green shades correspond to greater frequency of nominal (p<0.05 uncorrected) associations of the respective region with ^cis-eQTL^AGE-PRSs across the 25 PRS thresholds tested. D2 PRS associations are overlayed in red (pFDR significant associations), and yellow (pFDR trending). Abbreviated brain regions correspond to full names accordingly: PRC = precentral, PCU = precuneus, cMF = caudal middle frontal, IN = insula, SMG = supramarginal, TT = transverse temporal, ST = superior temporal, PAC = paracentral, SF = superior frontal, pTR = pars triangularis, mOF = medial orbitofrontal, MT = middle temporal, pOR = pars orbitalis, PCL = pericalcarine, PH = parahippocampal, rAC = rostral anterior cingulate, pOP = par opercularis.

#### 3.1.2 ^GWAS^AGE-PRS associations with cortical thickness

In contrast to the ^cis-eQTL^AGE-PRS results, no regional CT associations survived FDR correction within each of the 10 ^GWAS^AGE-PRSs. However, there was a distributed pattern of nominally significant associations. This consisted primarily of lower CT across the caudal anterior cingulate (n=4 ^GWAS^AGE-PRSs), postcentral (n=3 ^GWAS^AGE-PRSs), fusiform (n=1 ^GWAS^AGE-PRSs), superior frontal (n=1 ^GWAS^AGE-PRSs), and paracentral (n=1 ^GWAS^AGE-PRSs) gyri. Additionally, nominal associations of greater CT were present for the rostral anterior cingulate (n=2 ^GWAS^AGE-PRSs, pars opercularis (n=1 ^GWAS^AGE-PRSs) caudal middle frontal (n=1 ^GWAS^AGE-PRSs), pericalcarine (n=1 ^GWAS^AGE-PRSs) gyri. Figure 5 summarizes associations across all 9 ^GWAS^AGE-PRS and CT associations, and Figure 6 depicts regional ^GWAS^AGE-PRS and CT associations. Full statistical results can be found in Supplementary File 2.

**Figure 5.**
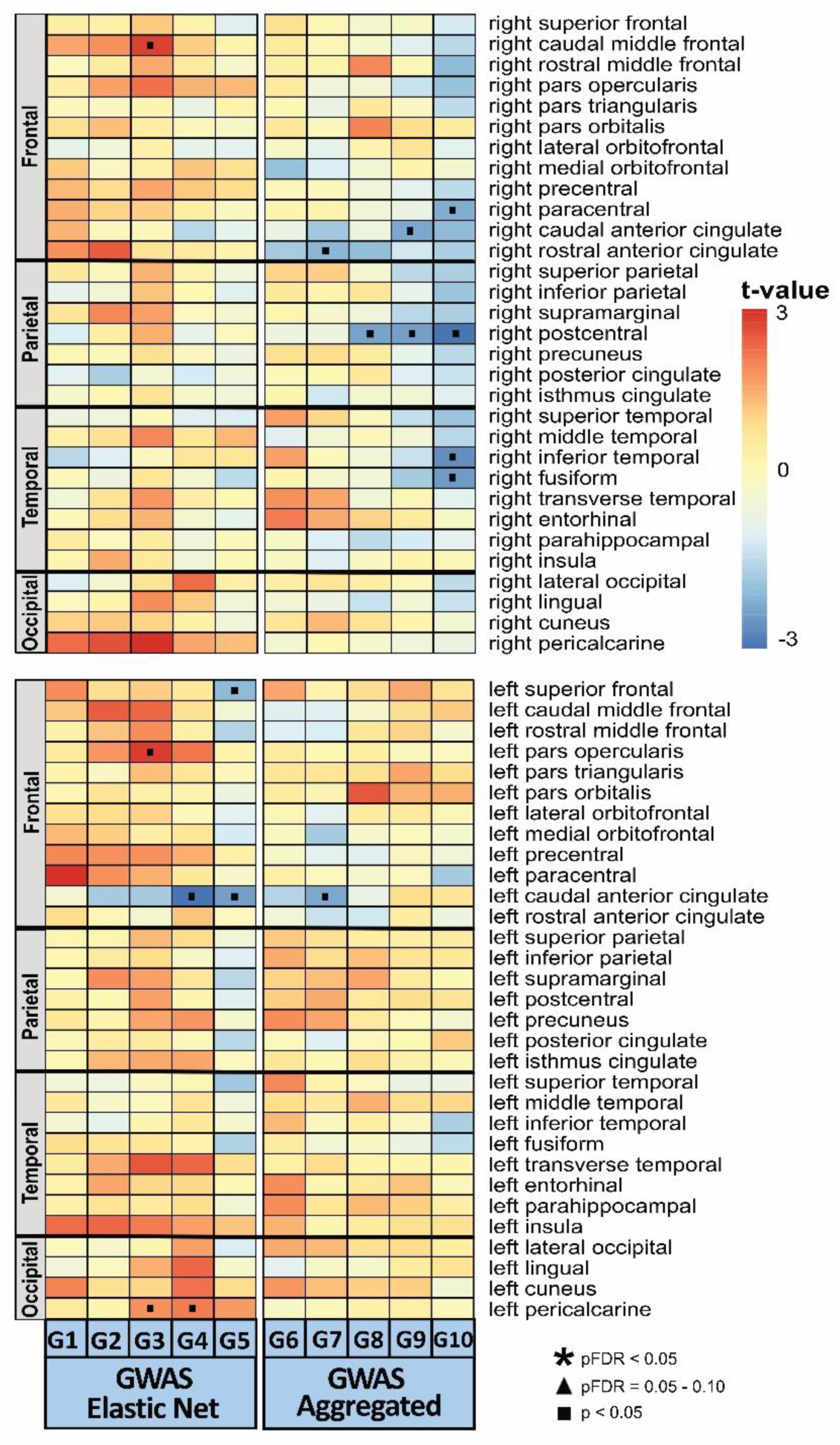
Overview of ^GWAS^AGE-PRSs associations with cortical thickness. Each column corresponds to one of the 10 ^GWAS^AGE-PRSs, while rows depict their respective associations with hemispheric brain regions (top = right hemisphere, bottom = left hemisphere). Within each hemisphere, regions are further organized by frontal, parietal, temporal, and occipital lobes. Warm colors (red/orange) reflect positive associations (i.e., higher PRS → greater cortical thickness), while cool colors (blue) reflect negative associations (i.e., higher PRS → lower cortical thickness) in the respective brain region. Asterisk symbols represent FDR-significant regional associations, while triangle symbols represent FDR-trending associations, and square symbols represent significant associations at p<0.05 uncorrected.

**Figure 6.**
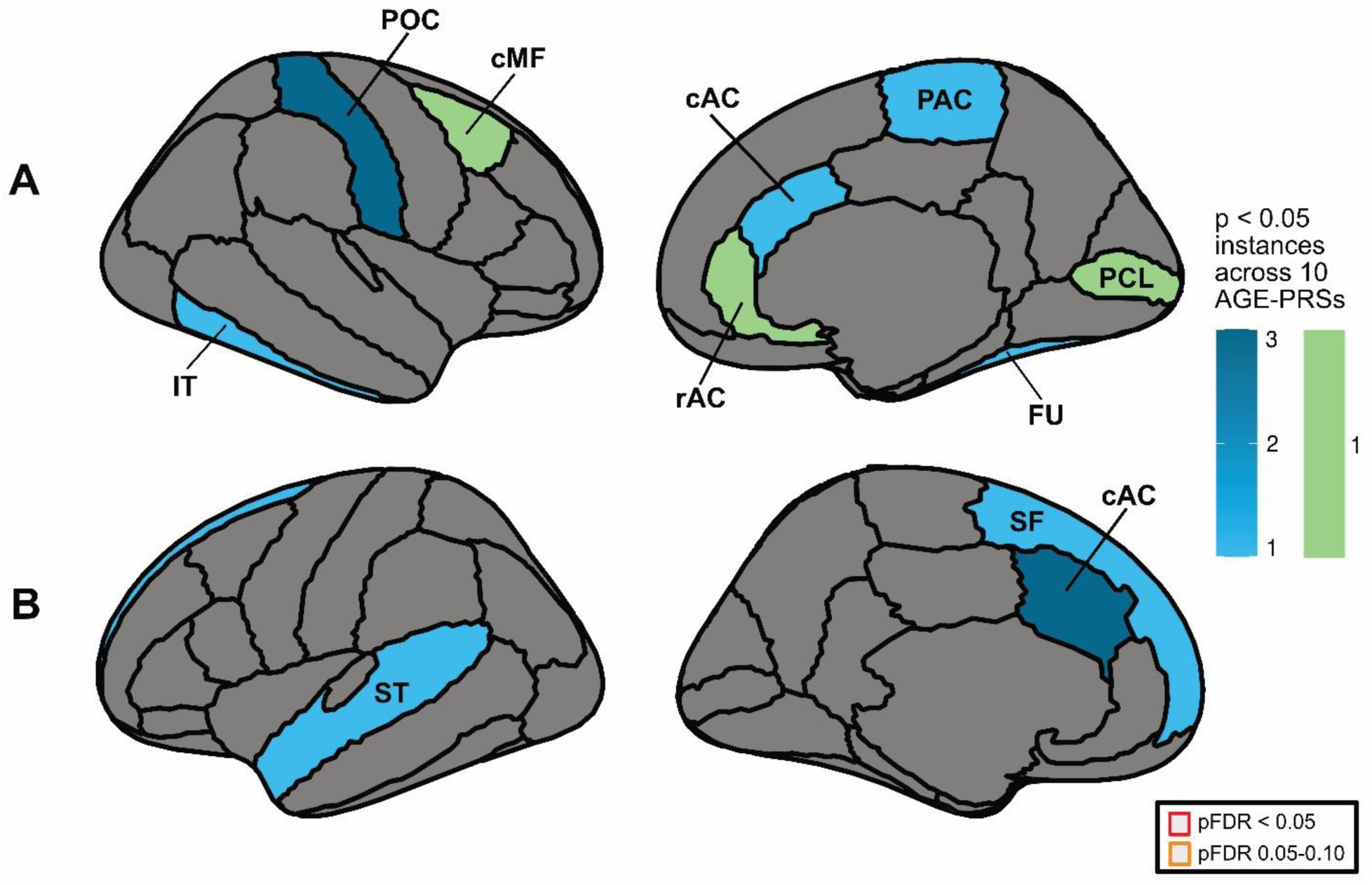
Regional brain map of ^GWAS^AGE-PRSs associations with cortical thickness across thresholds. Brain atlases depicting greater cortical thickness (green) and lower cortical thickness (blue) are displayed for the right hemisphere (A) and left hemisphere (B). Darker shades correspond to greater frequency of nominal (p<0.05 uncorrected) associations of the respective region with ^GWAS^AGE-PRSs across the 10 PRS thresholds tested. No pFDR significant (red) or pFDR trending (yellow) associations were present. Abbreviated brain regions correspond to full names accordingly: cAC = caudal anterior cingulate, POC = postcentral, SF = superior frontal, FU = fusiform, PAC = paracentral, IF = inferior temporal, SF = superior temporal, cMF = caudal middle frontal, PCL = pericalcarine, rAC = rostral anterior cingulate.

### 3.2 AGE-PRS associations with cognitive performance

#### 3.2.1 ^cis-eQTL^AGE-PRS associations with cognitive performance

Significant associations between ^cis-eQTL^AGE-PRS and better performance on the Trail Making Test B and Trail Making Test B-A tasks occurred in score D1 (q<0.01, 542 SNPs) (pFDR=0.035, pFDR=0.035). Furthermore, pFDR trending levels of this association were present in scores D2, E2, E1, C1, and C2, in descending order of magnitude. Statistical details of these trending associations can be found in Supplementary Table 3. Additionally, a significant association with worse performance on the Symbol Digit Substitution task occurred in score A5 (q<0.30, 371 SNPs) (pFDR=0.049), and at trending level in score A4 (q<0.20, 287 SNPs) (pFDR=0.056).

Lastly, a trending-level association with worse performance on Trail Making Test A was present in score A4 (q<0.20, 287 SNPs) (pFDR=0.056). No significant associations (all pFDR>0.238) were present between ^cis-eQTL^AGE-PRS and Tower Rearranging, Numeric Memory, and Fluid Intelligence tasks. Supplementary Figure 1 depicts ^cis-eQTL^AGE-PRS associations with cognitive performance. Full statistical results can be found in Supplementary Material 3.

#### 3.2.2 ^GWAS^AGE-PRS associations with cognitive performance

No significant associations occurred between ^GWAS^AGE-PRS and performance across the cognitive assessments, with all pFDR>0.264. Full statistical results can be found in Supplementary Material 4.

### 3.3 AGE-PRSs by sex interactions

No evidence of a significant interaction between ^cis-eQTL^AGE-PRS and regional CT was found (all pFDR>0.093). Similarly, no evidence of an interaction between ^GWAS^AGE-PRS and regional CT was present (all pFDR>0.186). Full statistical results can be found in Supplementary Material 5 and 6, respectively.

## 4. Discussion

Here, we provide evidence that genetic risk for accelerated molecular brain aging has an identifiable neurostructural and cognitive signature in mid- to late-life. Specifically, ^cis-eQTL^AGE-PRSs were associated with greater CT in age-sensitive frontal, temporal, and parietal regions, accompanied most robustly by better executive function performance on Trail Making Test B. Conversely, only nominal associations were present between ^GWAS^AGE-PRSs and CT, primarily with lower CT in age-sensitive frontotemporal regions. These results suggest that in non-clinical populations, genetic risk for accelerated molecular brain may be paradoxically associated with neurostructural and cognitive resilience in mid- to early old age. This resilience profile may reflect previously uncharacterized pathways of brain reserve or accelerated adaptation to aging, which may in turn inform novel strategies for prevention of age-related cognitive decline.

### 4.1 ^cis-eQTL^AGE-PRS associations with CT and cognition

In our main results, ^cis-eQTL^AGE-PRSs were associated with widespread greater CT, most robustly in the bilateral precentral and left insula, as well as the right supramarginal and precuneus regions. Further trending level associations occured in the bilateral caudal middle frontal and right insula regions. These frontal, temporal, and parietal regions display age-related thinning in normal aging, as do the nominal associations observed in the orbitofrontal cortex and temporal regions (Fjell et al., 2009; Salat et al., 2004; Thambisetty et al., 2010). Thus, while directionally opposite, our findings are consistent with the spatial distribution of age-related neurostructural effects. It is notable that patterns of lower CT also did occur considerably, but not at a significant level (Figure 3). Of note, while most regions showed bilateral effects, the precuneus was strongly lateralized, with 17 associations (16 nominal, 1 FDR significant) in the right hemisphere, and only one (nominal) in the left, suggesting the right precuneus may have particular involvement in aging and aging-related pathology, and warrants consideration in future studies.

The unexpected findings of greater CT are not necessarily contrary to brain aging-related changes. Most notably, this phenotype could indicate compensatory or resiliency processes in response to pathology precipitated by increased expression of age-related genes, consistent with the notion of brain “reserve”. In particular, brain reserve refers to variation in individual neurobiological capital supporting brain structure, facilitating better outcomes in brain aging pathology prior to clinical or cognitive changes (Stern et al., 2020), and could possibly reflect our greater CT findings. Further, brain reserve is thought to be supported by brain maintenance, the genetic and modifiable interplay of lifestyle factors supporting brain reserve (Stern, 2012).

While the molecular and cellular mechanisms underlying brain reserve and brain maintenance are not fully understood, increased neurotrophic factors, synaptogenesis, angiogenesis, and healthy glial functioning are thought to contribute (Serra and Gelfo, 2019).

Similarly, cognitive reserve refers to the adaptability and efficiency of cognitive processes in response to brain aging pathology (Stern, 2009). Cognitive reserve may account for the robust higher executive function performance (Trail Making Test B) observed across PRSs alongside concomitant findings of greater CT. It is notable that efficiency and interactions of functional brain networks are thought to underlie cognitive reserve (Stern et al., 2020), and our significant and trending frontoparietal regions partly overlap the two age-sensitive networks thought to subserve executive function (Hausman et al., 2021). Specifically, the frontoparietal network comprises large portions of the middle frontal gyrus (Yao et al., 2020), while the cingulo-opercular network contains both the insula and supramarginal gyrus regions (D’Andrea et al., 2023). Stronger connectivity in each of these networks is speculated to be a source of cognitive reserve, possibly subserved by thicker cortex (Stern et al., 2019). Furthermore, the precuneus is integral to the default mode network (DMN), wherein the rapid inhibition of activity necessary for executive function performance typically diminishes in aging (Brown et al., 2019). Greater CT in these particular regions may help delineate contributions to cognitive reserve and executive function adaptability in aging and aging-related pathology.

If the outcome of substantial greater regional CT is indeed driven by reserve mechanisms, the question remains as to which age-related pathologies the response is against. In our previous work (Lin et al., 2020), ^cis-eQTL^AGE-PRSs were associated with both diagnosis and lower cognitive functioning in both MDD and AD in clinical datasets. While MDD literature often finds patterns of cortical thinning associated with the disorder, there is also substantial literature of cortical thickening (Suh et al., 2019). In the present study, we observed regional overlap with MDD literature for greater CT in the precentral (Papmeyer et al., 2015; Zorlu et al., 2017), insula (Tu et al., 2012; Zorlu et al., 2017), supramarginal (Perlman et al., 2017; Qiu et al., 2014) and middle frontal gyri (Qiu et al., 2014) regions. Given the relatively limited literature on cortical thickening in MDD, it is unclear whether greater CT in these regions underlies pathology alone, reserve responses (Bansal et al., 2018; Zaremba et al., 2018), or a combination of each.

In addition to MDD, our prior evidence of associations between ^cis-eQTL^AGE-PRS and AD pathology (Lin et al., 2020) raises a particularly interesting parallel to the current findings. Generally, characteristic thinning in AD overlaps with our significant regions, namely the precuneus, supramarginal, superior and inferior frontal, and temporal regions (Dickerson et al., 2009). However, a recent biphasic model of AD has been proposed, whereby early AD pathology in late middle age manifests as initial widespread thickening of the cerebral cortex, alongside concomitant reduced mean diffusivity (Batzu et al., 2020; Montal et al., 2021; Williams et al., 2023). This phase precedes the eventual severe cortical atrophy and cognitive decline characteristic of the disorder in later life, and is thought to occur during high accumulation of amyloid beta prior to tau accumulation (Fortea et al., 2014). This thickening pattern has been observed in temporal and parietal regions mutual to our findings, but most notably in the precuneus, where we observed markedly greater CT in the right hemisphere (Fortea et al., 2010). While the mechanism is not fully known, biomarkers suggest the involvement of neural hypertrophy and activation of glial cells (Montal et al., 2018). Our findings of greater CT in a middle age to early old age population do not necessarily reflect AD pathology specifically, but this phenomenon of initial widespread cortical thickening in brain aging pathology may frame our findings of reserve as an initial response to pathology prior to, and predictive of, worsening outcomes in older age.

Speculatively, an alternative interpretation of the association between ^cis-eQTL^AGE-PRS and greater CT and executive function can be made through the lens of adaptive brain aging. From this perspective, age-related changes in gene expression reflect potentially anticipatory adaptations to the gradual and natural loss of physiological integrity associated with normal aging, rather than being pathological in and of themselves. Therefore, cis-eQTLs driving age-related changes in gene expression to occur earlier than expected would create a potentially beneficial phenotype akin to ‘accelerated adaptation’ to brain aging. Future work in longitudinal samples coupled with targeted molecular studies would be needed to test the viability of this interpretation.

### 4.2 ^GWAS^AGE-PRS associations with CT and cognition

Unlike ^cis-eQTL^AGE-PRSs, ^GWAS^AGE-PRSs reached only nominally significant associations with age-sensitive regions, mostly with lower CT. These included characteristic frontal regions (caudal anterior cingulate (cACC), superior frontal), temporal regions (fusiform, superior and inferior temporal) and parietal (paracentral) regions (Fjell et al., 2009; Lemaitre et al., 2012; Thambisetty et al., 2010). The lack of FDR-significant associations with CT and cognition was unexpected, given our hypotheses. This could be due, in part, to the age distribution of this middle age to early late age population (mean age of 64.1 years). It is possible the risk pathways captured by ^GWAS^AGE-PRSs are less impactful on CT thinning during middle-age, and that these associations may progress and become more pronounced in late-life.

Additionally, the GWAS performed to identify SNPs in scores G1:G5 was conducted in a discovery sample of 165 (Lin et al., 2020) and was not highly powered. Notably, most associations were with ^GWAS^AGE-PRSs G6:G10, where SNPs were aggregated from external datasets to increase quantity of SNPs. It is possible a higher powered GWAS identifying more SNPs associated with high delta brain age could yield a stronger signal. Of the nominal ^GWAS^AGE-PRSs associations, the cACC was most frequent across PRS thresholds. While findings of cACC thinning in normal aging sometimes conflict (Fjell et al., 2009; Salat et al., 2004; Thambisetty et al., 2010), pronounced cACC thinning is associated with AD (Jeong et al., 2021; Jones et al., 2006; Yang et al., 2019), suggesting ^GWAS^AGE-PRS may be associated with age-related pathology in this region.

The disparate phenotypic profiles associated with ^cis-eQTL^AGE-PRSs and ^GWAS^AGE-PRSs can be understood, in part, by the distinct and complementary nature of the SNPs included in each PRS family. Firstly, the GWAS captured associations with delta molecular brain age, a complex phenotype. The resulting SNPs may encompass wider, pleiotropic effects than those from the relatively straightforward cis-eQTL analysis. Furthermore, while we indexed cis-eQTLs variants nearby target genes, we could not identify trans-eQTLs influencing gene expression from further distances or different chromosomes, as their smaller effect size requires much larger populations (Kirsten et al., 2015). Since trans-eQTLs can act on multiple genes at once, and overlap with cis-eQTL target genes (Yao et al., 2017), their network of regulatory influence may have contributed additional unaccounted-for complex phenotypic effects. Notably, the SNPs comprising the separate PRSs across the two approaches were almost completely non-overlapping, with only two mutual SNPs being shared at the most permissive thresholds (G10: 394 SNPs, E5: 1838 SNPs). We also observed little spatial overlap in the regions significantly associated with each family of PRS. These non-overlapping effects highlight the value of modelling both types of genetic influence on brain aging, however, future studies should employ larger discovery samples to improve power for detecting GWAS-significant SNPs.

This current work is not without limitations. Firstly, the postmortem tissue in our previous work (Lin et al., 2020) was disproportionately derived from males (80%). This could create biased outcomes, as both regional changes and magnitude of thinning involve sex differences for this age range (Gautam et al., 2013; Thambisetty et al., 2010). However, after conducting exploratory analyses we found no significant sex interactions in our models.

Secondly, the available data from the UK Biobank was cross-sectional at a single time point. Ideally, future work with these PRSs could leverage longitudinal data extending into later age to both increase strength of inferences and reveal more pronounced effects of the PRSs. Additionally, we utilized tabular neuroimaging data in this study. Future work could benefit from employing vertex-wise data to capture more focal effects of the PRSs outcomes on CT. Lastly, due to the large scope of the UK Biobank, cognitive assessments are, by necessity, shortened to 5 minutes and self-administered. While the validity of Trail Making Test B is considered to be high (Fawns-Ritchie and Deary, 2020), the other age-sensitive assessments may not have captured cognitive outcomes of the PRSs in full. Future work would benefit from complete, well-validated assessments administered by a trained assessor.

Limitations aside, our work provides the first characterization of genetic risk for accelerated molecular brain aging on the structure of the living human brain. As such, it may grant valuable insight into regional correlates of structural brain reserve and cognitive reserve in age-related brain pathology, informing future translational research in identifying targetable pathways for treatment and prevention.

## Acknowledgments

YSN is supported by a Koerner New Scientist Award and a Paul Garfinkel New Investigator Catalyst Award administered by the CAMH Foundation, as well as a Discovery Grant from the Natural Sciences and Engineering Research Council of Canada.

This research has been conducted using the UK Biobank Resource under Application Number 85461.

---

**Supplemental Figure 1.**
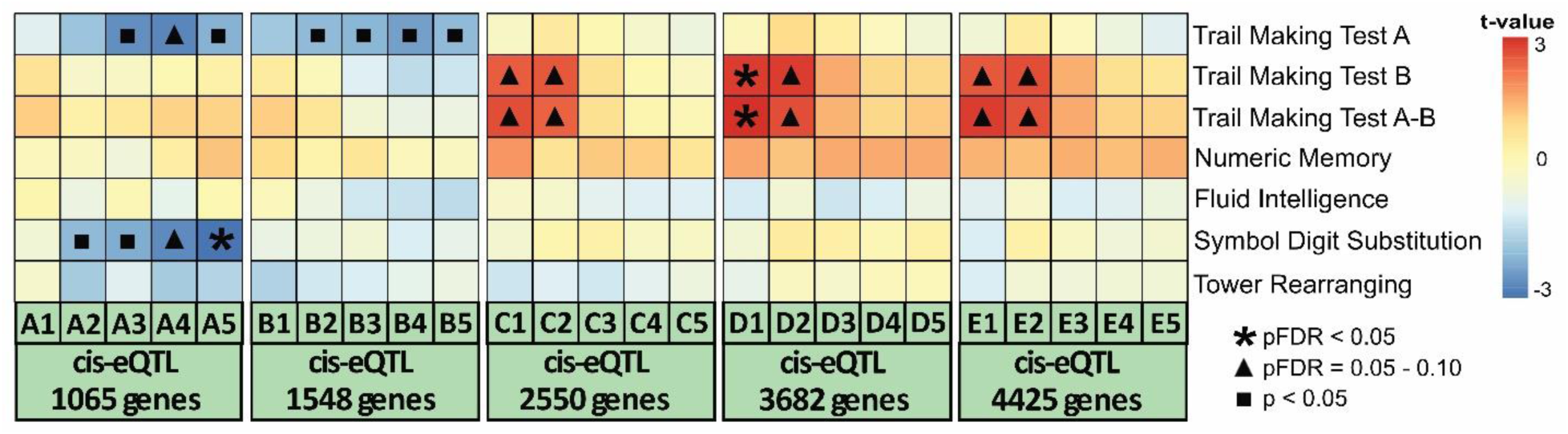
Overview of ^cis-eQTL^AGE-PRSs associations with cognitive performance. Each column corresponds to one of the 25 ^cis-eQTL^AGE-PRSs, while rows depict their respective associations cognitive test performance. Warm colors (red/orange) reflect positive associations (i.e., higher PRS → greater cortical thickness), while cool colors (blue) reflect negative associations (i.e., higher PRS → lower cortical thickness) in the respective brain region. Asterisk symbols represent FDR-significant regional associations, while triangle symbols represent FDR-trending associations, and square symbols represent significant associations at p<0.05 uncorrected.

